# Satellite repeat transcripts modulate heterochromatin condensates and safeguard chromosome stability in mouse embryonic stem cells

**DOI:** 10.1101/2020.06.08.139642

**Authors:** Clara Lopes Novo, Emily Wong, Colin Hockings, Chetan Poudel, Eleanor Sheekey, Simon Walker, Gabriele S. Kaminski Schierle, Geeta J. Narlikar, Peter J. Rugg-Gunn

**Affiliations:** Epigenetics Programme, Babraham Institute, Cambridge CB22 3AT, UK; Tommy’s National Miscarriage Research Centre at Imperial College London, W12 0NN, UK; Department of Biochemistry and Biophysics, University of California San Francisco, San Francisco, CA, USA; Department of Chemical Engineering and Biotechnology, University of Cambridge, Cambridge CB3 0AS, UK; Imaging Facility, The Babraham Institute, Babraham Research Campus, Cambridge CB22 3AT, UK; Wellcome Trust – Medical Research Council Cambridge Stem Cell Institute, University of Cambridge, Cambridge CB2 1QR, UK

**Keywords:** Heterochromatin, phase-separation, nuclear organisation, noncoding RNA, genome stability, embryonic stem cells, pluripotency

## Abstract

Heterochromatin maintains genome integrity and function, and is organised into distinct nuclear domains. Some of these domains are proposed to form by phase separation through the accumulation of HP1α. Mammalian heterochromatin contains noncoding major satellite repeats (MSR), which are highly transcribed in mouse embryonic stem cells (ESCs). Here, we report that MSR transcripts can drive the formation of HP1α droplets *in vitro*, and scaffold heterochromatin into dynamic condensates in ESCs, leading to the formation of large nuclear domains that are characteristic of pluripotent cells. Depleting MSR transcripts causes heterochromatin to transition into a more compact and static state. Unexpectedly, changing heterochromatin’s biophysical properties has severe consequences for ESCs, including chromosome instability and mitotic defects. These findings uncover an essential role for MSR transcripts in modulating the organisation and properties of heterochromatin to preserve genome stability. They also provide new insights into the processes that could regulate phase separation and the functional consequences of disrupting the properties of heterochromatin condensates.

## Introduction

Functional compartmentalisation of the genome segregates repetitive, gene-poor regions into constitutive heterochromatin^1^. Establishing and maintaining the appropriate regulation of heterochromatin is essential for preserving nuclear architecture, genome stability and DNA repair, and for silencing transposon expression^2,3^. In interphase cells, constitutive heterochromatin from different chromosomes cluster in cytologically defined nuclear bodies called chromocenters. These structures are characterised by the presence of histone H3 lysine 9 trimethylation (H3K9me3) and of heterochromatin protein 1α (HP1α)^4–6^. In the context of purified components, phosphorylation or DNA binding drives soluble HP1α into phase-separated droplets^7,8^. Furthermore, earlier studies showed that human HP1 proteins display dynamics on the order of seconds within heterochromatin puncta^9,10^. These and other observations in cells support the possibility that chromocenters could form by phase separation through the localised accumulation of HP1α. A phase-separation based model has important implications for how heterochromatin is assembled and regulated. However, the underlying biological processes that could promote HP1α-mediated phase separation *in vivo*, and the functional consequences of disrupting the phase separation of heterochromatin domains, remain as yet unknown.

The regulation of constitutive heterochromatin is developmentally controlled^11^ and this regulation is perturbed during cell stress and ageing^12^. Interestingly, in mouse, constitutive heterochromatin adopts an unusual configuration in both embryonic stem cells (ESCs) and early embryonic cells, compared to most other cell types, as it has reduced levels of H3K9me3, dynamically binds heterochromatin-associated proteins, and has an uncompacted chromatin structure^13–21^. Such distinct properties also extend to cytological differences, for example, constitutive heterochromatin is structured into much larger and fewer chromocenters in ESCs compared to most somatic cells^14,15,22,23^. How and why heterochromatin in ESCs adopts this unusual configuration is unknown, although one clue could be that ESCs transcribe high levels of noncoding major satellite repeat (MSR) elements, which constitute the predominant DNA sequences within chromocenters^13,15,19,24^ MSR transcripts remain close to their sites of transcription at chromocenters, and biochemical data indicate that MSR RNA can interact with heterochromatin-associated proteins, such as HP1α and SUV39H2^25–27^. Importantly, the control of MSR transcription is integrated into the transcription factor networks that sustain ESCs, which suggests that the regulation of heterochromatin in ESCs is an active process that is tightly controlled and coupled to cell state^15^. Evidence that MSR transcripts potentially have a functional role in ESCs comes from studies of early mouse development, which demonstrate that MSR RNA is required to establish chromocenter formation and for embryo development to proceed^28–30^. An important goal, therefore, is to understand the role that repetitive, noncoding RNAs, such as MSR transcripts, play in promoting the biophysical properties of heterochromatin formation and stability.

Here, we report that noncoding satellite RNA alters the physical properties and nuclear organisation of heterochromatin in mouse ESCs. We found that MSR transcripts catalyse a permissive and dynamic environment within individual chromocenters and promote the formation of large heterochromatin foci that are a hallmark of pluripotent cells. We further show that MSR RNA can drive the formation of HP1α droplets *in vitro* and is required for HP1α organisation within heterochromatin domains *in vivo*. The reduction of MSR transcripts prompted chromocenter properties to transition to a less dynamic and more stable heterochromatin state, and triggered the reorganisation of chromocenters into more numerous, smaller and compact foci. This aberrant chromocenter reorganisation led to rapid chromosome instability, as reflected by the appearance of chromosome end-fusions and pericentric gaps. These findings thus uncover an unexpected protective function for heterochromatin condensates in preserving genome stability.

## Results

### Constitutive heterochromatin forms liquid-like condensates in mouse embryonic stem cells

Studies of HP1α behaviour in *Drosophila* and in mammalian cells have proposed that heterochromatin is possibly held within liquid-like, phase-separated compartments^7,8^. To define the biophysical properties of constitutive heterochromatin in mouse ESCs, we used time-lapse imaging to track the movement and dynamics of individual chromocenters in live cells. For that, we generated a stable ESC line that expressed a doxycycline-inducible, monomeric GFP fusion protein that has a transcription activator-like effector (TALE) targeting the MSR sequence (Figures 1A and B)^31^. Fluorescent microscopy confirmed that the induced TALE-MSR-GFP signal localised to chromocenters throughout the cell-cycle (Supplementary Figure 1A and Supplementary Video 1), similar to previous observations^31^. As expected, the targeting of TALE-MSR-GFP to MSR elements did not affect chromocenter organisation in ESCs (their number and size, Supplementary Figures 1B and 1C, respectively). Importantly, our time-lapse imaging of TALE-MSR-GFP revealed that individual chromocenters separated and coalesced rapidly, with the movement of chromocenter extrusions occurring within minutes (Figure 1C and Supplementary Video 2). The ability to undergo such dynamic processes are features attributed to liquid-like membraneless condensates^32^. Liquid-like condensates are also characterised by the rapid, internal movement of molecules^33,34^. And so to further assess the dynamics of molecules within chromocenters of ESCs, we used fluorescence recovery after photo-bleaching (FRAP) to measure the mobile (liquid) and immobile (static) TALE-MSR-GFP-bound chromatin fractions. Because TALEs can bind with strong affinity (low nM Kd values) to their target DNA, both *in vitro* and *in vivo*^35,36^, this assay allows us to measure the physical properties of chromocenters, including both the movement of heterochromatin and the dynamics of chromatin-associated proteins. One chromocenter per nuclei was bleached, and the recovery of the GFP signal in this region over time quantified. The fluorescent signal recovered quickly at photo-bleached chromocenters, reaching 50% of the initial intensity within ~30s after bleaching (Figure 1D and Supplementary Video 3), with comparable dynamics to other molecules that were previously reported to be within liquid-like condensates^37–39^. In addition, an immobile, stable component of ~25% remained after photo-bleaching (Figure 1E), which is in line with measurements in other cell types^8,36^. The recovery could arise from the dissociation of bleached TALE-MSR-GFP molecules and their replacement with unbleached TALE-MSR-GFP molecules or from movement of the chromatin region itself. Compared to the time-scales that we observe, previous studies in ESCs have reported substantially slower recovery rates of bleached histones (t_1/2_ = ~100s) and faster recovery of soluble heterochromatin proteins, such as HP1α-GFP (t_1/2_ = ~3.5s;^14^). Thus, we favour the interpretation that we are measuring a combination of both replacement and movement. Based on these results, we conclude that DNA binding factors, such as TALE-MSR-GFP, bound to constitutive heterochromatin in ESCs display behaviours that are consistent with being part of a phase-separated compartment that consists of both dynamic and stable components.

**Figure 1:**
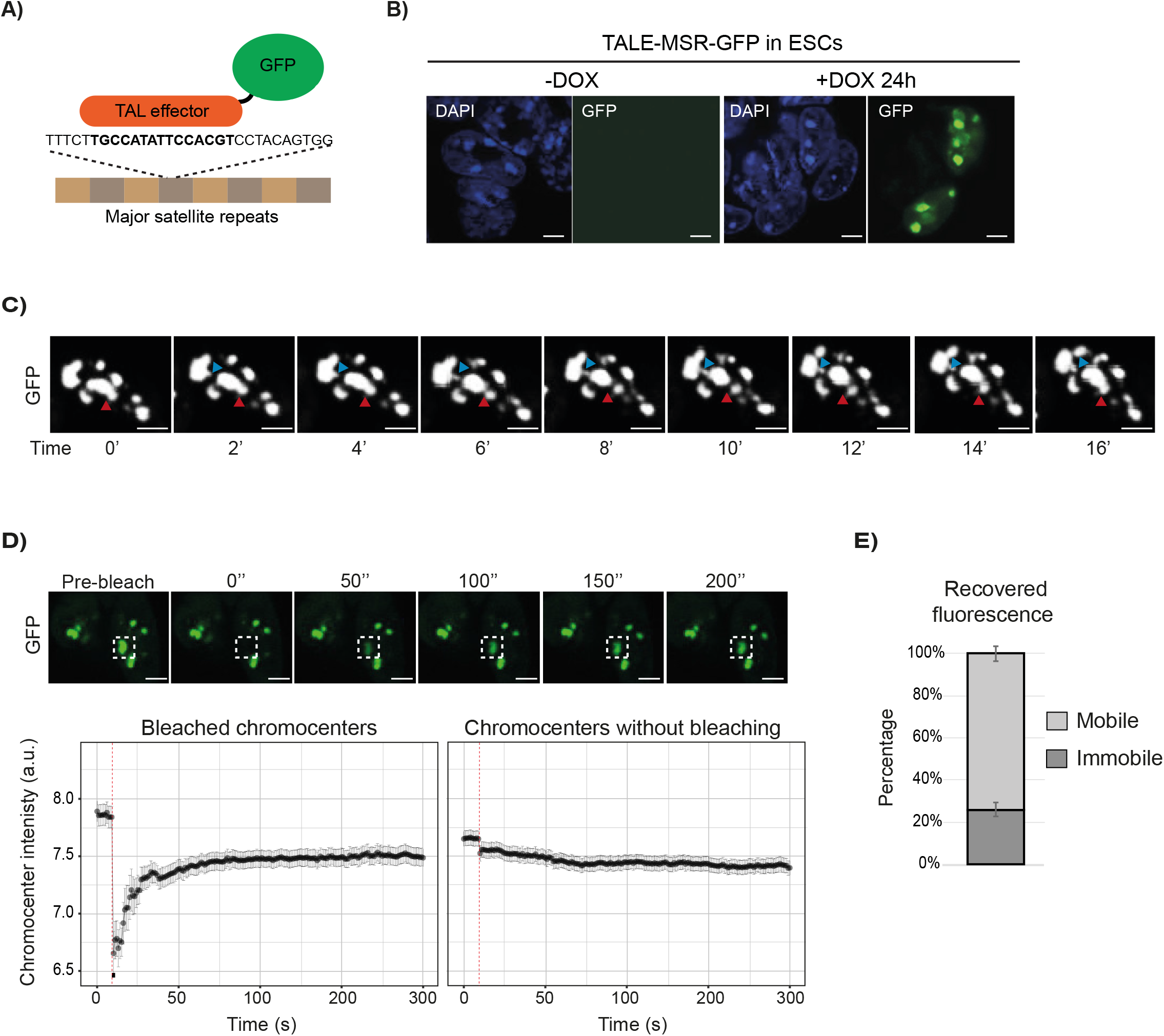
Constitutive heterochromatin forms liquid-like condensates in mouse embryonic stem cells. A) A schematic showing how doxycycline-inducible TALE-GFP is recruited to MSR DNA to enable live-cell visualisation of chromocenter dynamics. B) Representative images of nuclei showing the TALE-MSR-GFP signal and DAPI counterstain in ESCs without and with doxycycline induction. Scale bars, 5μm. C) Stills from a time-lapse microscopy experiment in doxycycline-induced TALE-MSR-GFP ESCs. These images of an ESC nucleus over time reveal the ability of GFP-labelled chromocenters to separate and coalesce (red arrow) and to form protrusions (blue arrow). Scale bars, 5μm. D) Quantification of fluorescence recovery after photo-bleaching individual chromocenters. Top panel shows representative images of the FRAP experiment; the dashed squares indicate the photo-bleached region. Lower panel shows the recovery of the GFP signal (log2 mean intensity ± SEM) over time at either bleached (left) or unbleached (right) chromocenters. Data were collected from at least three independent photo-bleaching experiments and show a summary of 49 photo-bleached chromocenters. Scale bars, 10μm. E) Histogram depicting the percentage of mobile and immobile components of chromocenters, as calculated from the FRAP data. See also Supplementary Figure 1 and Supplementary Videos 1, 2 and 3.

### Satellite transcripts regulate constitutive heterochromatin behaviour in embryonic stem cells

Chromocenters are distinctively large and less numerous in ESCs^14,22,23^, which suggests that an as-yet unknown component acts to promote the fusion of these liquid-like heterochromatin condensates. Because MSR RNA levels are substantially higher in ESCs relative to most other cell types^13,15,24^, and because pull-down experiments indicate that MSR transcripts can interact with HP1α^25,27^, we hypothesised that MSR transcripts might contribute to the formation of heterochromatin condensates. To directly test this hypothesis, we applied a commonly used approach for noncoding RNA studies; we depleted both strands of MSR transcripts using sequence-specific locked nucleic acid (LNA) DNA gapmers^28^. LNA DNA gapmers deplete target transcripts by acting at transcriptional^40^ and post-transcriptional steps^41^. After transfecting ESCs with LNA DNA gapmers that target the MSR transcripts (MSR gapmers), MSR RNA levels were reduced by ~50%, compared to cells transfected with control gapmers, as shown by RT-qPCR analysis (Figure 2A). The decreased level of MSR transcripts after LNA DNA gapmer transfection is comparable to that of MSR transcriptional levels in more differentiated cell types (Supplementary Figure 2A)^15^. The levels of other noncoding transcripts, such as of minor satellites or LINEs, were unchanged after MSR gapmer treatment (Figure 2A). In addition, we verified that the depletion of MSR RNA in ESCs did not affect the expression of undifferentiated cell markers or promote cell differentiation (Supplementary Figures 2B and 2C).

**Figure 2:**
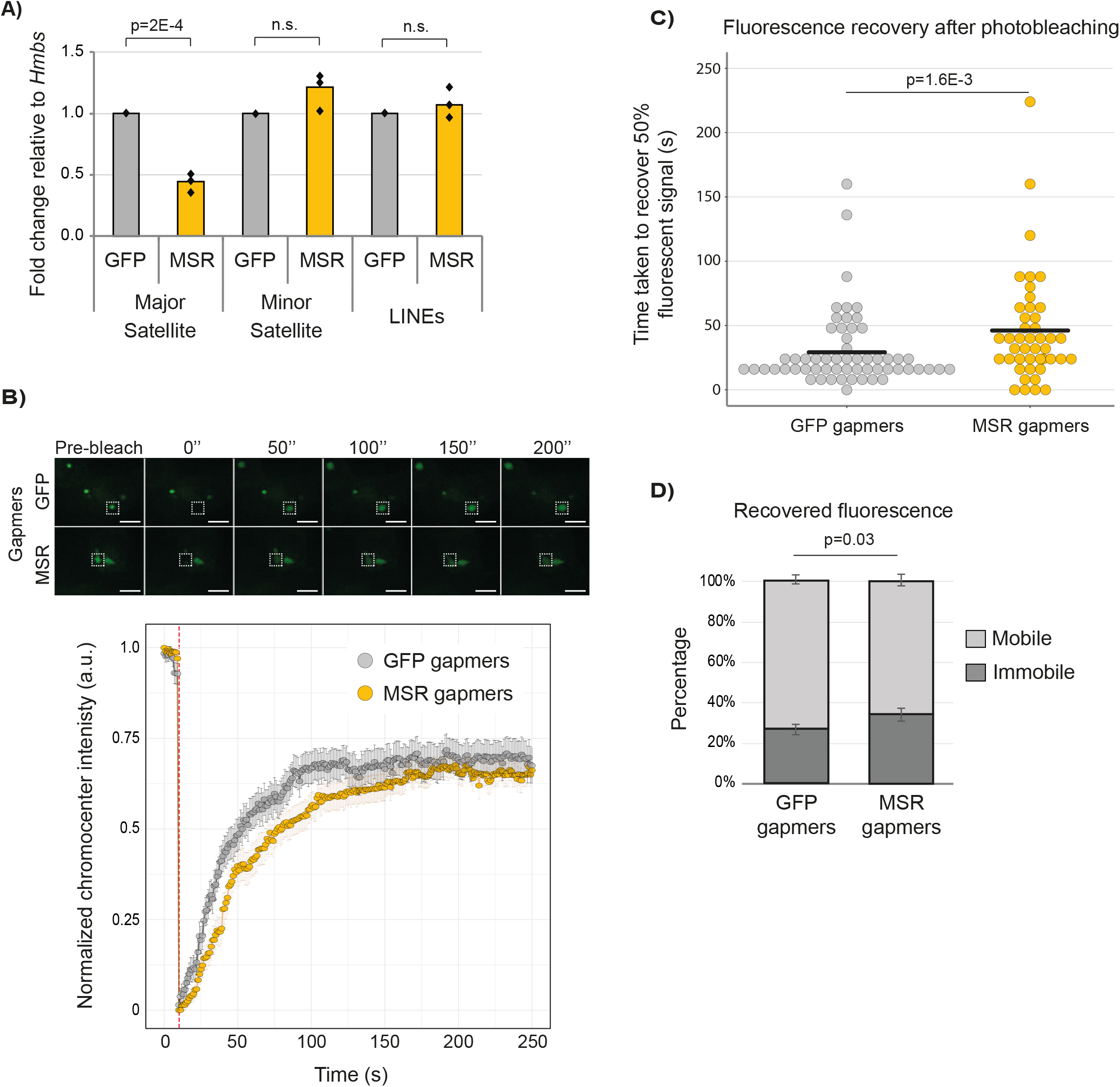
Major satellite repeat transcripts regulate constitutive heterochromatin behaviour in mouse embryonic stem cells. A) Bar chart showing depleted MSR transcript levels following the transfection of ESCs with LNA-DNA gapmers that target MSR but not control sequences (*GFP*). The transcripts of other repetitive sequences, such as minor satellites and LINE RNA, were not affected by the LNA-DNA gapmers. Individual data points show the mean values for three independent experiments and were compared using an unpaired, two-sided t-test. B) Top panel shows representative images of timelapse FRAP experiments in GFP or MSR gapmer-transfected ESCs. The dashed squares indicate the photo-bleached areas at 0 s; note that the size of the bleached area was the same in all experiments. The lower panel shows GFP intensity (normalised mean intensity ± SEM) of photo-bleached chromocenters (red dotted line shows moment of photo-bleaching) over time after GFP or MSR gapmer transfection. Data were collected from five independent photo-bleaching experiments. Scale bars, 10μm. C) Chart showing the increased time taken to recover 50% of the TALE-MSR-GFP signal at bleached chromocenters in ESCs with depleted MSR RNA levels following MSR-targeting gapmers. Each circle represents one photo-bleached chromocenter. Data collected from five independent experiments (n=49 for GFP and n=45 for MSR) and were compared using an unpaired two-sample Mann-Whitney test. D) Histogram depicting the mobile and immobile components of chromocenters, as calculated from the FRAP data. Data were collected from at least five independent photo-bleaching experiments (n=49 for GFP and n=45 for MSR) and the mobile/immobile proportions between the two experiments were compared using an unpaired two-sample Mann-Whitney test. See also Supplementary Figure 2 and Supplementary Video 4.

We next used live-cell imaging of TALE-MSR-GFP to measure chromocenter dynamics in ESCs after depleting MSR transcripts. We confirmed that following the depletion of MSR transcripts, TALE-MSR-GFP still localised to chromocenters, as expected (Supplementary Figure 2D). Excitingly, FRAP experiments revealed that the fluorescence recovery time at photo-bleached chromocenters doubled in MSR RNA-depleted cells compared to controls, indicating that the dynamic movement of TALE-MSR-GFP molecules within chromocenters was reduced (Figures 2B, 2C and Supplementary Video 4). Notably, the recovery immediately after photo-bleaching was substantially slower in the MSR RNA depleted cells (Figure 2B) and this reduction in recovery is consistent with the TALE proteins being bound within a less dynamic chromatin^14^. Furthermore, the proportion of mobile GFP signal decreased slightly in the MSR RNA depleted cells (Figure 2D), suggesting that the combination of dynamic and stable components within chromocenters is also affected by MSR RNA levels. Together, these data lead us to propose that MSR RNA catalyses a dynamic, physical environment within individual chromocenters.

### Satellite RNA promotes phase-separation of HP1α

We next investigated a potential mechanism by which MSR RNA might contribute to the biophysical properties of heterochromatin. As MSR RNA binds HP1α^25,27^, we hypothesised that MSR RNA might affect the ability of HP1α to phase-separate. Excitingly, adding *in vitro* transcribed MSR RNA (272 nucleotides containing a single repeat sequence) to recombinantly purified human HP1α induced droplet formation in an HP1α concentration-dependent manner (Figure 3A). Furthermore, MSR RNAs containing multiple repeat sequences promoted droplet formation at progressively lower HP1α concentrations, comparable to, and in some cases better than, enzymatically synthesised polyuridylic acid (polyU) at identical nucleotide concentrations (Figure 3A and Supplementary Figure 3A). While both strands of MSR RNA exhibited the ability to form droplets, transcripts corresponding to the forward strand consistently yielded larger droplets as compared to the reverse strand (Figure 3A and Supplementary Figure 3A). These results suggest that MSR RNAs are capable of binding to HP1α and promote thematically similar behaviour as has been observed for HP1α with DNA *in vitro*^7,8^. This model provides a molecular basis for understanding why HP1α-associated heterochromatin disperses into smaller condensates after MSR RNA levels are depleted (Figure 3B). And, conversely, it offers an explanation as to why the induction of MSR transcription in differentiated cell types can drive the fusion of chromocenters into larger aggregates^15^.

We next used structured illumination microscopy (SIM) to test whether a reduction in MSR RNA levels affects HP1α structure at chromocenters. Upon MSR transcript depletion, HP1α formed a smaller and more compact core at chromocenters, concomitant with a stronger overlap between HP1α and H3K9me3 marked volumes (Figures 3B, 3C and Supplementary Figure 3B). Consistent with this finding, we also detected increased HP1α signal at chromocenters by immunofluorescence microscopy (Figure 3D) and at the underlying satellite DNA by chromatin immunoprecipitation (ChIP) (Figure 3E), following MSR RNA depletion. We next investigated whether the increase of chromatin-associated HP1α at chromocenters affected chromatin compaction by using SiR-DNA fluorescence lifetime methodology that measures chromatin compaction levels in defined genomic compartments^42^. Following MSR RNA depletion, we observed using this approach more compacted chromatin was present in heterochromatin domains but not in euchromatic regions of the nucleus (Figure 3F). Together, these results lead us to conclude that high levels of MSR RNA in ESCs promote the fusion and aggregation of chromocenters into large foci, and also prevent premature heterochromatinisation and compaction of chromatin within these domains.

**Figure 3:**
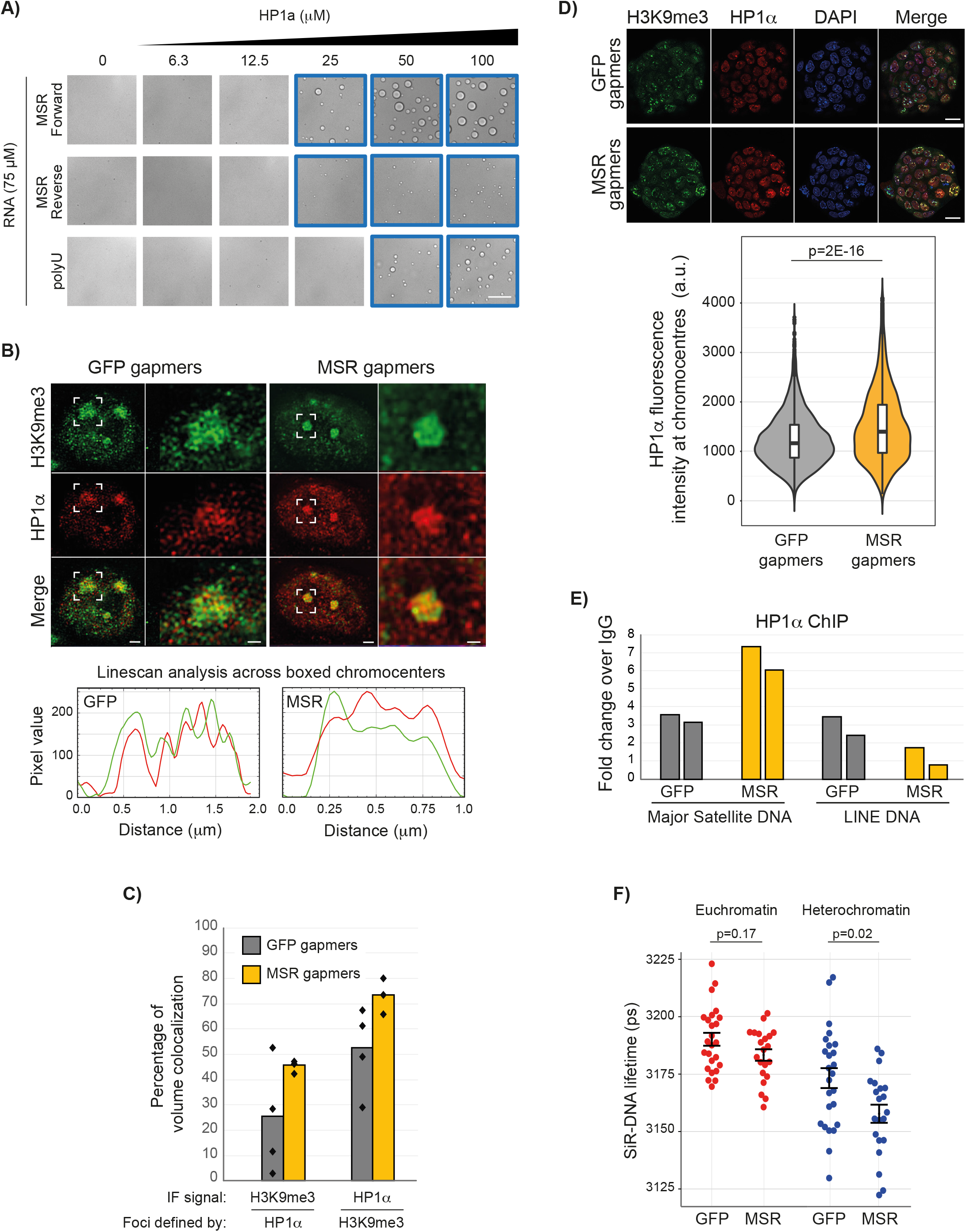
Satellite RNA promotes phase-separation of the heterochromatin protein, HP1α. A) *In vitro* transcribed MSR RNAs, or synthetic polyU RNA, were added to increasing concentrations of recombinant HP1α protein. Images framed in blue show the conditions that allowed for the formation of HP1α droplets. All images show the same magnification; scale bar, 100μm. B) Representative images of 3D reconstructions acquired with structured illumination microscopy (SIM), showing details of H3K9me3 and HP1α organisation at chromocenters in ESCs that were transfected with GFP or MSR gapmers (scale bars, 2μm). White boxes indicate zoomed-in areas shown to the right (scale bars, 0.5μm). Linescan analyses of the boxed chromocenters are shown below and reveal the overlap of heterochromatin marks at chromocenters (HP1α, red line; H3K9me3, green line). C) Bar chart showing the percentage of H3K9me3 and HP1α 3D volumes that colocalise with either HP1α- or H3K9me3-marked foci following the GFP or MSR gapmer transfection of ESCs. Foci volumes were defined from at least three 3D-reconstructions of SIM for each gapmer condition (individual data points are shown). D) Upper, representative images of immunofluorescence microscopy showing H3K9me3, HP1α and DAPI signals upon gapmer transfection of ESCs with GFP or MSR gapmers. Scale bars, 20μm. Lower panel, quantification of HP1α immunofluorescence intensity at chromocenters that were defined by DAPI. A total of 2849 and 3209 chromocenters were quantified after two independent GFP and MSR gapmer transfection experiments, respectively, and were compared with a two-sample Mann-Whitney test. E) Histograms of ChIP results show the fold-change over IgG of HP1α bound at MSR DNA or LINE DNA upon transfection of ESCs with GFP or MSR gapmers. Two biological replicates are shown side by side. F) Plot shows the fluorescence lifetime of the far-red dye SiR-DNA in both euchromatin and heterochromatin regions of nuclei following the transfection of ESCs with GFP or MSR gapmers. Each point represents the mean lifetime intensity per image from three independent experiments, each with >8 images per sample. Samples were compared with a two-sample Mann-Whitney test; error bars show mean ± SEM. See also Supplementary Figure 3.

### Satellite RNA scaffolds heterochromatin organisation in embryonic stem cells

Having established the effect of MSR RNA in promoting rapid chromatin dynamics within constitutive heterochromatin, we next investigated the role of MSR transcripts in chromocenter organisation. Line-scan microscopy analysis revealed that MSR RNA depletion triggered the rapid reorganisation of heterochromatin into more numerous, brighter and smaller chromocenters (Figures 4A-C), an organisational pattern that is characteristic of differentiated cell types. In addition, immunofluorescence microscopy and ChIP revealed the increased accumulation of H3K9me3 at heterochromatin domains after MSR RNA depletion (Figures 4D and 4E), relative to controls. These results further support that changes occur to the state of heterochromatin at chromocenters upon MSR RNA depletion. We conclude, therefore, that MSR transcripts are required to maintain the highly dynamic and distinctive organisation of heterochromatin domains in ESCs.

**Figure 4:**
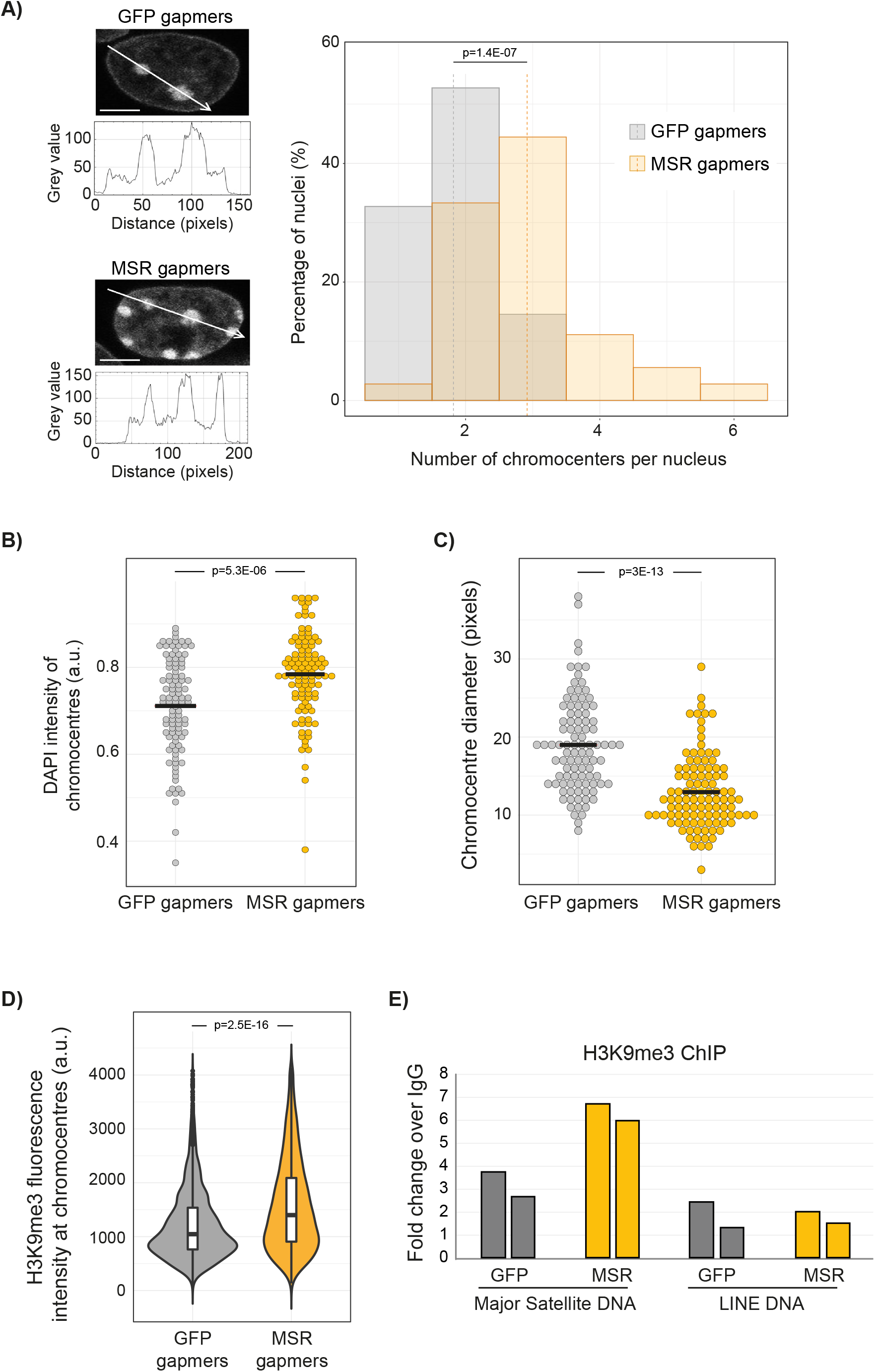
Degradation of satellite transcripts induces chromatin remodeling in embryonic stem cells. A) Left: Chromocenter organisation as revealed by microscopy and linescan analysis of DAPI signal from a single stack (2D) after ESCs were transfected with GFP or MSR gapmers. Scale bars, 5μm. Histogram (right) shows the number of chromocenters per nucleus. Data were collected from three independent experiments and compared using a two-sample Mann-Whitney test. B, C) Plots show DAPI intensity (B) and diameter (C) of chromocenters per nucleus. A total of 100 and 105 chromocenters were quantified following the transfection of ESCs with GFP and MSR gapmers, respectively. Data were collected from three independent experiments and compared using a two-sample Mann-Whitney test. (D) Quantification of H3K9me3 immunofluorescence intensity at chromocenters that were defined by DAPI. A total of 2849 and 3209 chromocenters were quantified following two independent GFP and MSR gapmer transfection experiments, respectively, and were compared with a two-sample Mann-Whitney test. E) Histograms of ChIP results show the fold-change over IgG of the H3K9me3 signal at MSR DNA or LINE DNA following GFP or MSR gapmer ESC transfections. Two biological replicates are shown side by side.

### Chromocenter architecture protects chromosome stability in embryonic stem cells

Disrupted heterochromatin maintenance is often associated with the onset of chromosome instability, elevated DNA damage, and with defective mitosis. In somatic cells, this type of heterochromatin perturbation is typically triggered by the weakening of heterochromatin-associated processes^3^. ESCs are unusual, however, in that they can seemingly tolerate a permissive and uncompacted heterochromatin state without adverse consequences^43,44^. We therefore investigated whether a distinct heterochromatin state is not only tolerated in ESCs but perhaps also required to maintain proper chromosome segregation. Interestingly, we observed-shortly after MSR RNA depletion - that a large proportion of metaphase-stage nuclei showed clear hallmarks of chromosome instability (Figures 5A and 5B). Specifically, ~20% of nuclei had chromosome fusions, ~4% contained chromosome breaks, and ~30% showed signs of fragility at pericentromeric regions; defects that were rarely observed in control samples. Furthermore, immunofluorescence microscopy revealed a significant increase in the number of γH2AX foci in chromocenters after gapmer treatment (Figure 5C), which indicates the presence of elevated damage at satellite DNA following MSR RNA depletion.

**Figure 5:**
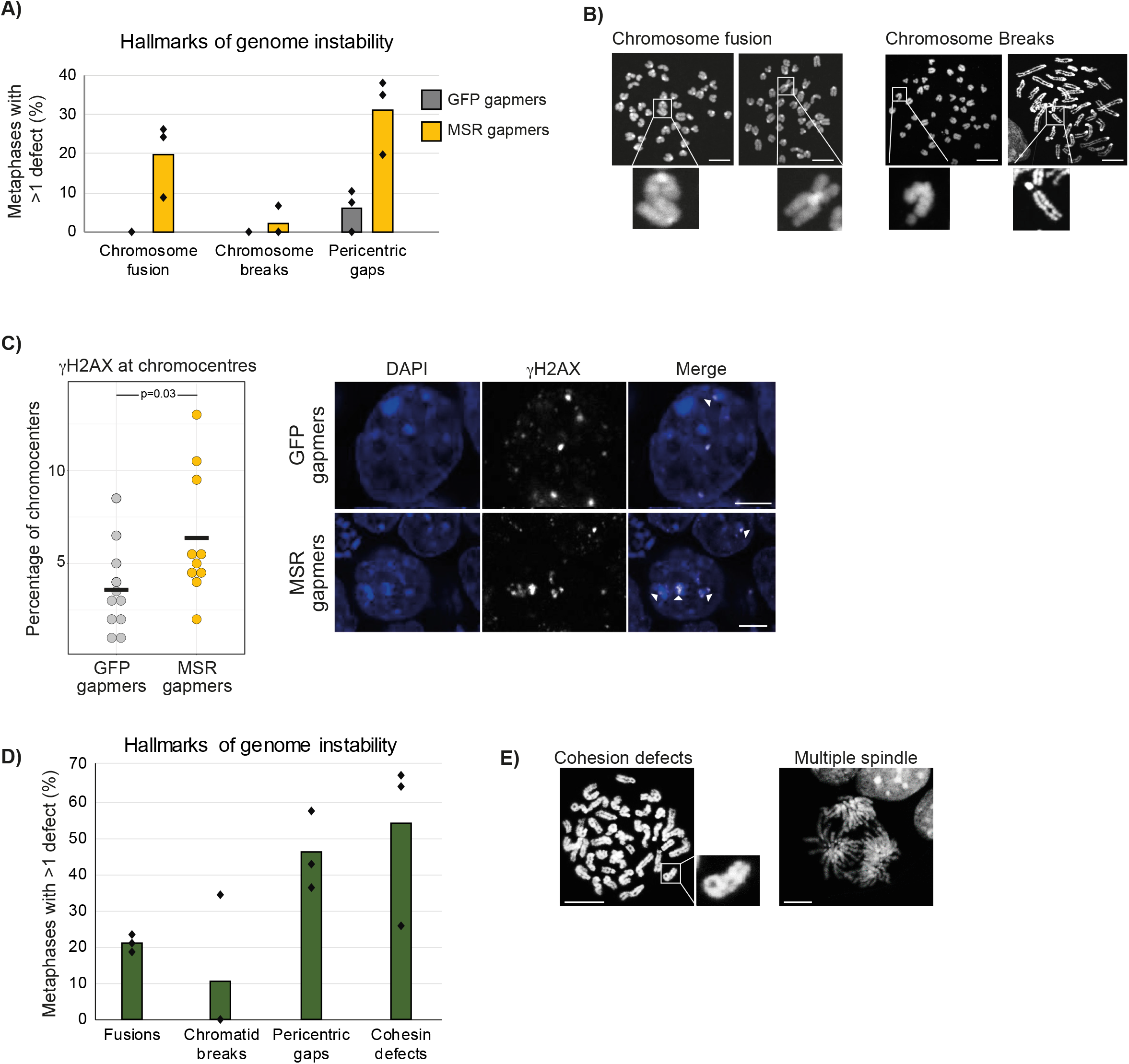
Chromocenter architecture protects chromosome stability in embryonic stem cells. A) Histogram showing that ESCs transfected with LNA-DNA gapmers that target MSR transcripts have a high proportion of metaphases with hallmarks of genetic instability, such as chromosome end-fusions and pericentric gaps. At least 31 metaphases from three independent gapmer transfections. Data points show percentages for each biological replicate. B) Representative images of DAPI-stained metaphases, exemplifying the cytogenetic defects observed. Scale bars, 10μm. C) Left: histogram showing the percentage of gamma-H2AX foci at chromocenters. Each point represents the percentage in each image analysed. Right, representative images of gamma-H2AX at chromocenters (white arrow). Scale bars, 5μm. A total of 317 and 420 ESCs were scored for GFP and MSR gapmer transfection, respectively, and the two samples were compared with an unpaired Mann-Whitney test. D) Histogram showing the genetic defects observed after TALE-MSR-KRAB induction in ESCs. At least 30 metaphases from three independent replicates were scored. Data points show percentages for each replicate. E) Representative images of DAPI-stained metaphases exemplifying the cytogenetic defects observed after TALE-MSR-KRAB induction. Scale bars, 5μm.

We then tested this association using a different system, this time by forcing an increase in chromocenter heterochromatinisation using a doxycycline-inducible TALE-MSR-KRAB protein^45^. Consistent with our above observations, the activation of TALE-MSR-KRAB in ESCs substantially impaired chromosome segregation, with ~20% of metaphase-stage nuclei containing fused chromosomes, ~10% with chromatid breaks, ~50% with pericentromeric gaps and >50% of analysed mitotic chromosomes showing defects in sister-chromatid cohesion (Figures 5D and 5E). In contrast, samples taken from the same ESC line but without doxycycline induction were normal. These results establish that heterochromatin organisation plays an important protective role in ESCs to safeguard chromosomal stability.

## Discussion

Our study demonstrates that noncoding satellite RNA modulates the physical properties and nuclear organisation of heterochromatin. Satellite repeat elements are highly transcribed in ESCs, but it remains unknown whether such transcripts are simply a by-product of the permissive chromatin environment that defines pluripotent cells or whether they themselves fulfil a functional role in genome regulation^46^. We now demonstrate that satellite repeat transcripts are required to maintain constitutive heterochromatin in a more dynamic state that organises into large chromocenters *in vivo*. At the molecular level, modelling and biochemical studies have proposed that RNA molecules can promote liquid-liquid phase-separation by lowering the concentration required for proteins to phase separate and/or to modulate the physical properties of phase-separated condensates^33,47^. Additionally, RNA molecules themselves can phase-separate into either liquid- or gel-like states, forming compartments that can sequester proteins and other RNAs^37,48^. Although noncoding transcripts have diverse sequences in different species, there are likely to be universal and sequence-independent principles for how RNA molecules promote the formation of nuclear domains.

Our findings show that depleting MSR RNA disrupts the dynamics of heterochromatin, increases heterochromatinisation and alters chromocenter organisation. These results also provide an interpretation for a previous observation that an unknown nuclear RNA component can sustain the higher-order structure of pericentric heterochromatin^49^, and presents evidence that the ability to maintain chromocenter stability resides at least in part with the transcript itself. As satellite repeats are under the transcriptional control of pluripotency factors in ESCs^15^, these new findings provide a direct connection between cell state and heterochromatin regulation. Thus, a key developmental switch is controlled by the decline in pluripotency factor availability as cells differentiate, which triggers the downregulation of MSR transcripts and the reorganisation of chromocenters towards a configuration that is typical of somatic cells. Mechanistically, we show that MSR RNA is a ligand that promotes the ability of HP1α to phase separate *in vitro*, likely by modulating a similar closed-to-open equilibrium as has been hypothesised to be modulated by DNA^7^. General modeling of protein-nucleic acid condensation suggests that the addition of nucleic acids reduces the thermodynamic and kinetic barriers of phase separation^50^, consistent with our observations here. An important implication arising from our work is that MSR RNA levels can be titrated through their transcriptional control and by their degradation and clearance. This could enable the fine tuning of heterochromatin properties into states that range from soluble to liquid to gel-like, perhaps to regulate chromatin accessibility, genome compartmentalisation and/or chromosome structure. An important future direction for research will be to investigate how this pathway integrates with other processes, including protein phosphorylation and alternative ligands, to direct heterochromatin properties and dynamics^51^.

In addition to scaffolding HP1α, our work raises the possibility that MSR transcripts have a second key function: to prevent the untimely and deleterious compaction of chromatin within heterochromatin domains. It remains unexplained how heterochromatin in ESCs remains permissive and uncompacted, despite residing within H3K9me3 and HP1α marked territories^16,17,52,53^. We propose that high levels of MSR RNA prevent HP1α from binding to satellite DNA, thereby sequestering HP1α away from chromatin and unable to compact it. This model is supported by biochemical experiments, which have shown that HP1α has a stronger preference for RNA than DNA^27^. As RNA and DNA bind to the same hinge region on HP1α, MSR RNA can effectively out-compete satellite DNA for binding to HP1α^27,54,55^. A reduction in MSR RNA levels would, therefore, enable HP1α to engage with and compact chromatin. We tested this prediction and found that the depletion of MSR RNA triggered the increased binding of HP1α at satellite DNA and increased chromatin compaction within chromocenters. These data also agree with previous findings that the transcriptional downregulation of MSR RNA upon ESC differentiation is closely coupled to the progressive compaction of chromatin^15,19,56^ and, conversely, that activating MSR transcription in fibroblasts results in the decompaction of chromocenters^15,19,56^.

Taken together, we propose that our findings support a model in which heterochromatin-associated MSR transcripts act near their sites of synthesis to hold HP1α molecules in close proximity to chromatin, but prevents full engagement and compaction. This increase leads to the formation of large condensates and keeps HP1α away from chromatin, in order to sustain a permissive heterochromatin structure in ESCs (Figure 6). Upon MSR RNA depletion, either experimentally or after ESC differentiation, HP1α is then released and can bind to satellite DNA, leading to the observed changes in heterochromatin compaction and organisation. It remains important to determine whether a reduction in MSR RNA concentration within heterochromatin condensates can trigger specific transitions in the material properties of heterochromatin condensates, with parallels to enhancer condensates^57^, or whether the relocalisation of HP1α and other processes is also required. Interestingly, a recent study showed that HP1 molecules rapidly exchange in and out of chromocenters in fully differentiated mouse cells^56^. Such rapid exchange is consistent with the low surface tension of phase-separated condensates that arise from electrostatically driven multi-valent interactions, such as those between DNA and HP1^51,58^. Excitingly, the results of this study raise the potential for a dosedependent effect of other charged ligands of HP1, such as MSR transcripts, in regulating heterochromatin by modulating the ability of HP1α to form multivalent interactions. These interactions perhaps transition chromocenters from being liquid-like aggregates, such as those described in early embryonic development^8^ and in pluripotent stem cells (as reported in this study), to being ‘ordered collapsed globules’ in differentiated cells^56^. It remains important to define whether HP1α alone could be the chromatin-binding factor that bridges and collapses chromatin into globules or if other factors, possibly cell-specific, are also involved.

**Figure 6:**
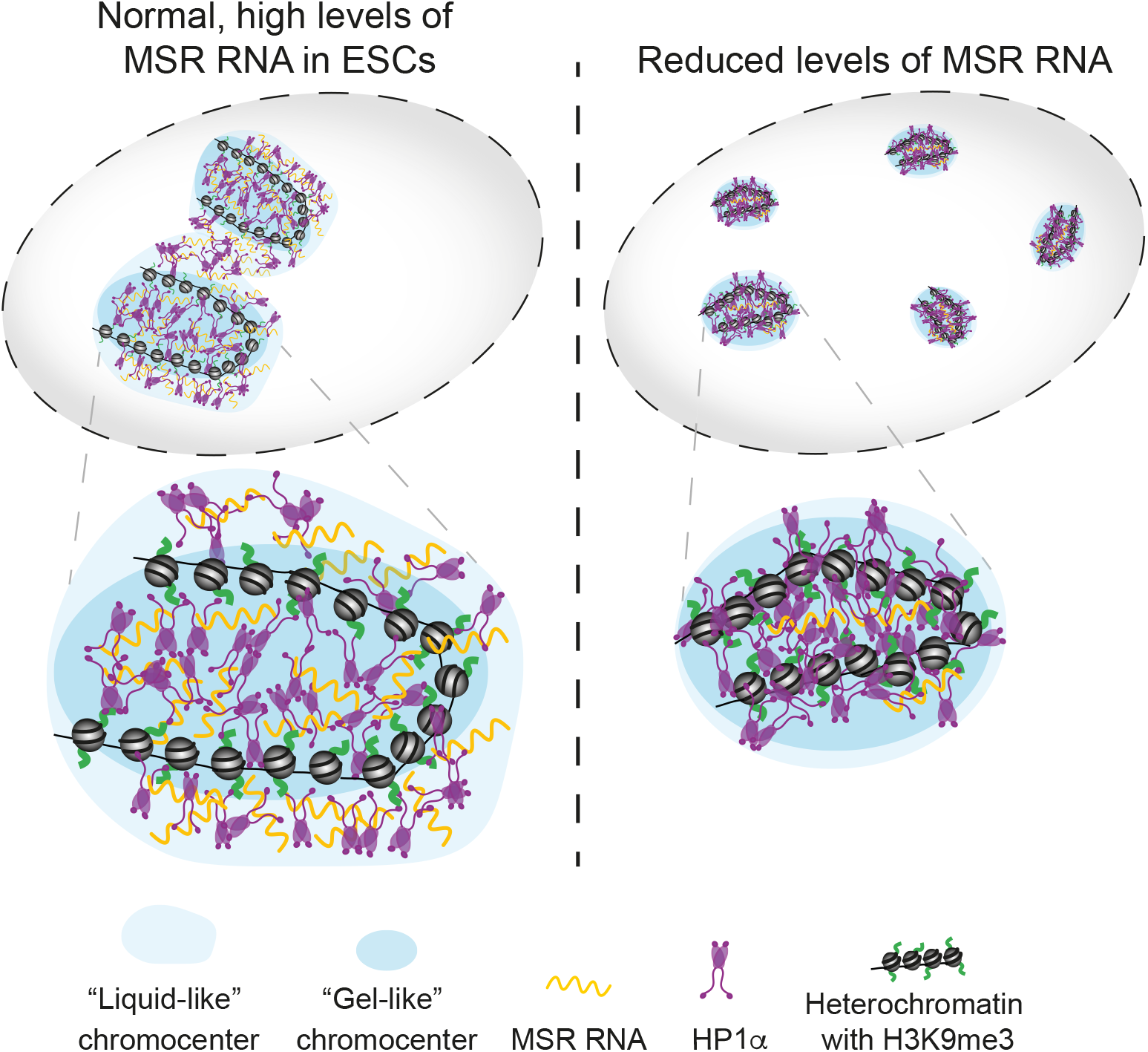
Model of the proposed role for MSR transcripts in chromocenter organisation. MSR transcripts contribute to chromocenter organisation, by: i) catalysing weak interactions between HP1α molecules; and ii) preventing HP1α from binding to chromatin to sustain a permissive heterochromatin structure in ESCs. Upon MSR RNA depletion, which normally occurs during ESC differentiation, the substrate for weak interactions is reduced and more HP1α molecules bind to satellite DNA, leading to the observed changes in heterochromatin compaction and organisation.

Pericentromeric heterochromatin forms one of the subdomains of centromeres, and preserving heterochromatin stability, including HP1α function is required for appropriate chromosome condensation and segregation in most cell types^59–63^. Curiously, ESCs can tolerate mutations that weaken heterochromatin pathways, while the same perturbation in somatic cells causes significant karyotypic defects^43,44^. Unexpectedly, we found in ESCs that MSR transcript depletion and strengthened heterochromatinisation in ESCs led to the rapid appearance of mitotic defects, highlighting the functional differences of heterochromatin regulation that exists between pluripotent and somatic cells. It is currently unclear why ESCs require this unusual form of heterochromatin regulation to maintain chromosome stability. One possibility is that this requirement arose concomitantly with the very short G1 phase of the cell cycle in ESCs^64,65^, perhaps to accommodate the rapid reassembly of chromocenters before the start of the next replication round. Changes to the biophysical properties of chromocenters could perturb the correct timing of DNA replication and/or transcription, which could quickly lead to the observed genetic instability defects. More generally and importantly, our findings imply that an association exists between the biophysical state of heterochromatin and the preservation of chromosome stability. Notably, noncoding transcripts are accumulated in other processes in which the genome is remodelled, such as in the formation of senescence-associated heterochromatin foci, cellular stress, embryo development and in human ageing and disease^29,66–70^. Based on the findings reported here, further work is needed in these other contexts to investigate the role of RNA molecules in controlling heterochromatin properties and genome function.

## Online Methods

Additional reagent and resource details are provided in Supplementary Table 1.

### Cell culture

Male, E14Tg2a mouse ESCs (129P2/OlaHsdl passages 19-28)^71^ were cultured in ESC media containing DMEM supplemented with 15% FBS, 1mM sodium pyruvate, 0.1mM 2-mercaptoethanol, O.1mM nonessential amino acids, 2mM Glutamax and 1000 U/ml LIF. Cells were maintained at 37°C on mitotically inactivated mouse embryonic fibroblasts and transferred for two passages onto gelatin-coated surfaces prior to collection. ESCs were routinely verified as being mycoplasma-free using a PCR-based assay (Sigma). The E14Tg2a line is not on the list of commonly misidentified cell lines (International Cell Line Authentication Committee).

Stable, doxycycline-inducible ESC lines were made by transfecting E14Tg2a cells using Lipofectamine 2000 (ThermoFisher) with 1ug PB-TET-TALE-MSR-mClover-ires-mCherry^15^ or with PB-TET-TALE-MSR-KRAB-ires-GFP plasmids, together with 1ug pCAG-*rtTA-Puro* and 2ug pCyL43 piggyBac transposase, followed by selection with 1.2ug/ml puromycin. Some ESC lines were flow sorted to purity based on mCherry or GFP expression following transient doxycycline induction, and were expanded for >10 passages in the absence of doxycycline. Doxycycline was applied at 0.5-1ug/ml for 24h to induce transgene expression.

### Depletion of major satellite repeat transcripts

ESCs were grown to 70% confluency on gelatin-coated plates and were transfected with LNA gapmer oligos (Exiqon) at a concentration of 100nM, using Lipofectamine RNAiMaX Reagent (ThermoFisher), following the manufacturer’s instructions. To ensure the robust depletion of target transcripts, the LNA gapmer transfections were repeated at 48h and 72h after the first transfection. Cells were collected for further analysis 24h after the final transfection. The LNA gapmer sequences are: LNA DNA gapmer GFP (gagaAAGTGTGACAagtg), LNA DNA gapmer Major Satellite 1 (acatCCACTTGACGActtg) and LNA DNA gapmer Major Satellite 2 (tattTCACGTCCTAAagtg), where the lowercase letters denote the LNA nucleotides^28^.

### RT-qPCR

For most RT-qPCR experiments, RNA was extracted using the RNeasy Mini Kit (Qiagen), reverse transcribed using Superscript II (Life Technologies) and random primers (Promega), and subjected to qPCR analysis, as previously described^72^. To analyse major satellite and other repeat classes, total RNA was extracted using TRIzol (Life Technologies) and treated with two rounds of 1U DNase I (Fermentas) per 1μg RNA in the presence of RiboLock RNase inhibitor (Fermentas) to remove genomic DNA. RNA (1μg) was reverse transcribed using Superscript II (Life Technologies) and random primers (Promega), in the presence of RiboLock RNase inhibitor. cDNA was amplified with SYBR Green Jumpstart Taq Ready Mix (Sigma) using primers from^73,74^ Primer sequenced; Major Satellite For, GACGACTTGAAAAATGACGAAATC; Major Satellite Rev, CATATTCCAGGTCCTTCAGTGTGC; Minor Satellite For, TGATATACACTGTTCTACAAATCCCGTTTC; Minor Satellite Rev, ATCAATGAGTTACAATGAGAAACATGGAAA; LINE L1 For, CTGGCGAGGATGTGGAGAA; LINE L1 Rev, CCTGCAATCCCACCAACAAT; Nanog For, ATGCCTGCAGTTTTTCATCC; Nanog Rev, GAGGCAGGTCTTCAGAGGAA; Klf4 For, ACACTTGTGACTATGCAGGCTGTG; Klf4 Rev, TCCCAGTCACAGTGGTAAGGTTTC; Hmbs For, CGTGGGAACCAGCTCTCTGA; Hmbs Rev, GAGGCGGGTGTTGAGGTTTC; T For, TCAGCAAAGTCAAACTCACCAACA; T Rev, CCGAGGTTCATACTTATGCAAGGA.

### Chromatin immunoprecipitation

Crosslinked ChIP experiments were performed as previously described^75^. Briefly, cells were fixed with 2mM DSG (Sigma) for 45min and then with 1% formaldehyde for 12min. Sonicated chromatin (250μg; 200-500bp fragments) was pre-cleared with blocked beads for 2h at 4°C and incubated at 4°C overnight with either 5μg HP1α antibody (ab77256, Abcam) or 1μg rabbit IgG (Jackson ImmunoResearch). Chromatin-antibody complexes were incubated for 6-8 hours at 4°C with protein A/G magnetic Dynabeads (Life Technologies), washed and the crosslinks were reversed. Native ChIP experiments for H3K9me3 (ab8898 Abcam) were performed as previously described^72^. ChIP DNA was analysed by qPCR using primers from^73,74^.

### In vitro transcription of major satellite repeats

MegaScript T3 and T7 kits (ThermoScientific) were used to transcribe forward and reverse transcripts of differing lengths from major satellite repeat DNA, according to the manufacturer’s instructions. To obtain transcripts with one single consensus repeat, oligos containing the forward or the reverse major satellite consensus sequences were synthesised with the T7 or the T3 sequences, respectively (GenScript) and hybridised. Two repeats were transcribed using the pCR4 Maj9-2 template (^73^; a gift from Thomas Jenuwein), which was digested with SpeI or NotI for T7 or T3 Megascript kits, respectively. Eight repeats were transcribed from the pySat template (^76^; a gift from from Niall Dillon, Addgene plasmid # 39238), which was digested with NotI or SalI for T7 or T3 Megascript kits, respectively. Prior to in vitro transcription, linearised plasmids were cleaned with 3M Sodium acetate and 100% ethanol. Linear templates (1μg) were in vitro transcribed according to manufacturer’s instructions, and RNAs were clean by LiCl precipitation.

### HP1α droplet-formation assay

Polyuridylic acid (polyU, Santa Cruz Biotechnology, cat no. sc-215733A) was dissolved in MilliQ water, then dialysed extensively against additional MilliQ water in a 3 kDa cutoff dialysis membrane (SpectraPor) to remove co-purified salts, ethanol precipitated and washed prior to final resuspension in MilliQ water. *In vitro* transcribed MSR RNAs and commercial polyU were quantitated using the total base hydrolysis method of RNA degradation^77^ and UV-absorbance quantitation at 260 nm^78^ to yield concentration in units of nucleotides. All RNAs were diluted in assay buffer (20 mM HEPES, pH 7.2, 75 mM KCl, 1 mM DTT) prior to use. Recombinant human HP1α protein was purified from *E. coli* as previously described^7^. Prior to use in droplet assays, protein was dialysed in the assay buffer (20 mM HEPES, pH 7.2, 75 mM KCl, 1 mM DTT).

Droplet assays were performed using a 384 Greiner Sensoplate (no. 781892) according to the protocol by Keenen and colleagues^79^ with a Nikon TiEclipse microscope fitted with a 20x DIC objective. Briefly, 2x solutions of HP1α (two-fold serial dilutions from 200 μM down to 1.6 μM, and 0 μM) were mixed in equal volume with a 2x solutions of RNA (150 μM nucleotide). The HP1α-RNA mixture was then vigorously pipet-mixed before transferring into the 384 well plate. Mixtures were allowed to settle for an hour prior to droplet imaging.

### Microscopy imaging and analysis

ESCs were cultured on glass coverslips precoated with 0.1% gelatin. For immunofluorescence experiments, cells were fixed with 3% paraformaldehyde in PBS for 10 min at room temperature (RT), washed three times with PBS for 5min, permeabilised with 0.1% Triton X-100 in PBS for 10 min, and washed three times with PBS for 5min. Cells were incubated with primary antibody (H3K9me3, 39161 Active Motif and HP1α, ab77256 Abcam) in blocking buffer (5% milk in PBS) and incubated either for 1h at RT or overnight at 4°C. They were then washed three times with PBS for 5min and incubated with secondary antibodies for two hours at RT. Images were collected on an Olympus FV1000 or Nikon A1-R confocal microscope. Coverslips were mounted onto slides with media containing DAPI (Vectashield H-1200, Vector Laboratories) and were imaged using a Nikon A1-R confocal microscope and a 60X oil objective. When indicated, a Z-stack of images was collected with 0.2 μm spacing and projected using maximum intensity. Image files were prepared using Fiji software. ImageJ software was used to quantify H3K9me3 foci, size and intensity using the ‘Analyze Particles’ tool (Novo et al., 2016). Linescan analysis was performed as previously described^17^. Colocalisation analysis was performed in Imaris (BitPlane, Oxford).

Super-resolution structured illumination microscopy images were acquired with a Nikon dual mode super-resolution microscope with structured illumination (SIM). The resulting reconstructed images were analysed in 3D in Imaris (BitPlane, Oxford).

### Live-cell imaging

TALE-MSR-GFP ESCs were grown on glass coverslips coated with 0.1% gelatin. Cells were imaged using an Andor Revolution spinning disk confocal microscope, which was equipped with a thermostatically controlled stage maintained at 37°C with a 60X oil immersion objective. A z-stack of images was collected with 0.5 μm spacing and projected using maximum intensity. Three biological replicates per cell line and doxycycline concentration were included in each experiment. At least 500 cells were analysed for each experimental group.

### Fluorescence recovery after photo-bleaching

FRAP experiments were performed on an Andor Revolution spinning disk confocal microscope. Images were acquired every second for 300 seconds (310 frames). The first ten frames were collected before the bleach pulse for baseline fluorescence. A 10×10 region surrounding a single chromocenter per cell was selected for bleaching puncta using 2% laser power (488 nm). Fluorescent intensities and image analysis were performed using the TrackMate plugin (v3.8.0) in the Fiji software^80^. After normalisation of the intensity signal, FRAP curves were generated from the fluorescence intensity of each chromocenter, at each timepoint (every second). To calculate the half-time [ln(2)/K] of fluorescence recovery after photo-bleaching, data was fitted to a one-phase association equation [Y=Y0 + (Plateau-Y0)*(1-exp(-K*x))], where Y0 is the fluorescence after photo-bleaching (GraphPad software). To correct for loss of fluorescence due to the bleaching pulse the mobile fraction was calculated as [Fm= (Plateau - First post-bleach) / (Pre-bleach - First post-bleach)] and the immobile fraction was measured as remaining fluorescence intensity unrecovered at plateau phase. FRAP experiments were repeated five times and data were acquired from one bleached and from at least one unbleached chromocenter per cell.

### Chromocenter organisation and compaction analysis

DAPI linescan analyses were performed using ImageJ on optical sections, where DAPI foci were at optimal focal planes. Fluorescence intensity histograms were generated with a linescan across the nucleus, and the background (outside of the nucleus) was subtracted from the baseline fluorescence within the nucleus and the chromocenter signal. Differences in chromocenter intensity are shown as the ratio of chromocenter peak height to nucleoplasmic signal.

For SiR-DNA-based analysis of chromatin compaction by FLIM^42^, cells were plated in 35 mm glass bottom dishes (P35-1.5-14-C, Mattek Corporation, MA, USA) the day before imaging. Cells were stained with 1 μM SiR-DNA (Spirochrome Ltd., Stein am Rhein, Switzerland) and with 10 μM verapamil (Spirochrome Ltd.) in cell culture medium for 1 hour, before changing to cell culture medium that contained 10 μM verapamil and 20 mM HEPES pH 7.4 for imaging. Fluorescence lifetime imaging (FLIM) was performed on a home-built confocal platform (Olympus Fluoview FV300), which was integrated with time-correlated single photon counting (TCSPC) to measure fluorescence lifetime in every image pixel. Output from a pulsed supercontinuum source (WL-SC-400-15, Fianium Ltd., UK, repetition rate 40MHz) was filtered using a bandpass filter FF01-635/18 to excite SiR-DNA. Fluorescence emission from the sample was filtered using 700/70nm (Comar Optics, UK) before passing onto a photomultiplier tube (PMC-100, Becker & Hickl GmbH, Berlin, Germany). Photons were recorded in time-tagged, time-resolved mode that permits the sorting of photons from each pixel into a histogram according to their arrival times. The data was recorded by a TCSPC module (SPC-830, Becker and Hickl GmBH). Photons were acquired for two minutes to make a single 256 × 256 FLIM image. Photon count rates were always kept below 1% of the laser repetition rate to prevent pulse pile-up. Photobleaching was verified to be negligible during acquisition. Approximately 10 representative images with several nuclei per field of view were acquired for each condition. FLIM images were analysed using FLIMfit v4.12.1 and fitted with a monoexponential decay function. Whole nuclei were segmented based on intensity (debris and mitotic nuclei were manually removed). A two-level mask that separated heterochromatin spots and euchromatin was created with Icy software spot detection tool^81,82^. Pixels in these two chromatin regions were binned and fitted separately to obtain two lifetime values for each image. Statistical analysis was carried out using an unpaired two-samples Mann-Whitney test.

### Quantification and statistical analysis

Statistical parameters including the exact value of n, SD, SEM and statistical significance are reported in the figures and the figure legends. For the majority of experiments, statistical significance is determined by the value of p < 0.05 by unpaired two-samples Wilcoxon test (Mann-Whitney test).

### Data availability statement

The data that support the findings of this study are available from the corresponding authors upon reasonable request.

### Code availability statement

No custom code was used.

## Acknowledgements

We are grateful to Niall Dillon, Maria Elena Torres-Padilla, Thomas Jenuwein and the Wellcome Trust Sanger Institute for kindly providing plasmids. We thank several Babraham Institute Facilities for their assistance, including the Flow Core, the Bioinformatics Group, and particularly Simon Andrews and Anne Segonds-Pichon, and also Hanneke Okkenhaug in the Imaging Facility. We thank members of P.J.R.-G.’s group for helpful discussions, Gavin Kelsey and Wolf Reik for comments on the manuscript, and the Babraham Institute Science Policy and Oversight Committee for their support. C.N. and P.J.R.-G. were funded by grants from the Biotechnology and Biological Sciences Research Council (BBS/E/B/000C0421, BBS/E/B/000C0422, BB/M022285/1 and the Core Capability Grant), and European Commission Network of Excellence EpiGeneSys (HEALTH-F4-2010-257082). E.W. was supported by an F32 fellowship from the NIH. C.H., C.P. and G.S.K.S. were supported by the Wellcome Trust, Alzheimer’s Research UK, the Michael J Fox Foundation, Infinitus China Ltd, and by the European Union’s Horizon 2020 Research and Innovation Programme under grant agreement number 722380. E.S. was supported by a British Society for Developmental Biology Gurdon / The Company of Biologists Summer Studentship. G.J.N. was supported by an MIRA grant from the NIH (R35 GM127020). C.N. is grateful to the Tommys National Miscarriage Centre for funding support.

## Author Contributions

Conceptualisation, C.L.N. and P.J.R-G.; Methodology, C.L.N., C.H., E.W., C.P., S.W.; Investigation, C.L.N., E.W., C.H., E.S.; Visualisation, C.L.N.; Writing – Original Draft, C.L.N. and P.J.R-G.; Writing – Review & Editing, C.L.N., E.W., G.J.N. and P.J.R-G.; Funding Acquisition, C.L.N., G.S.K.S., G.J.N., P.J.R-G.; Supervision, G.S.K.S., G.J.N. and P.J.R-G.

## Competing Interests

The authors declare no competing financial interests.

## Supplementary Figures

**Supplementary Figure 1:**
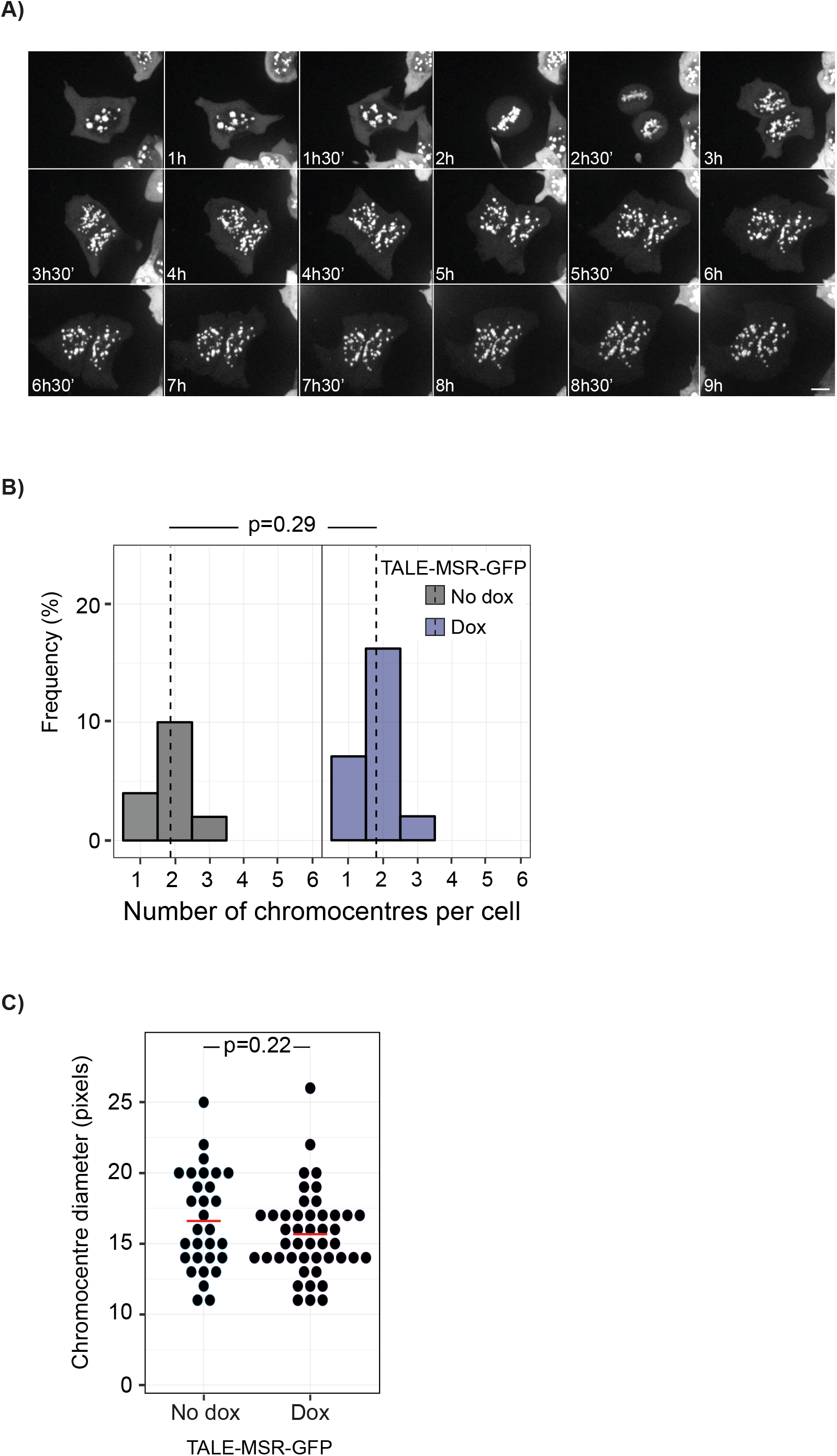
Inducible targeting of GFP to satellite DNA with a TALE does not affect the nuclear organisation of chromocenters, Related to Figure 1. A) Representative images of ESC nuclei from a time-lapse experiment, which show that TALE-MSR-GFP binds to major satellite sequences throughout the cell cycle. Scale bar, 5μm. B, C) Plots showing the number (B) and diameter (C) of chromocenter foci per ESC nucleus upon 24h doxycycline induction of TALE-MSR-GFP. Data were collected from three independent experiments and compared using a two-sample Wilcoxon test. See also Supplementary Videos 1–3.

**Supplementary Figure 2:**
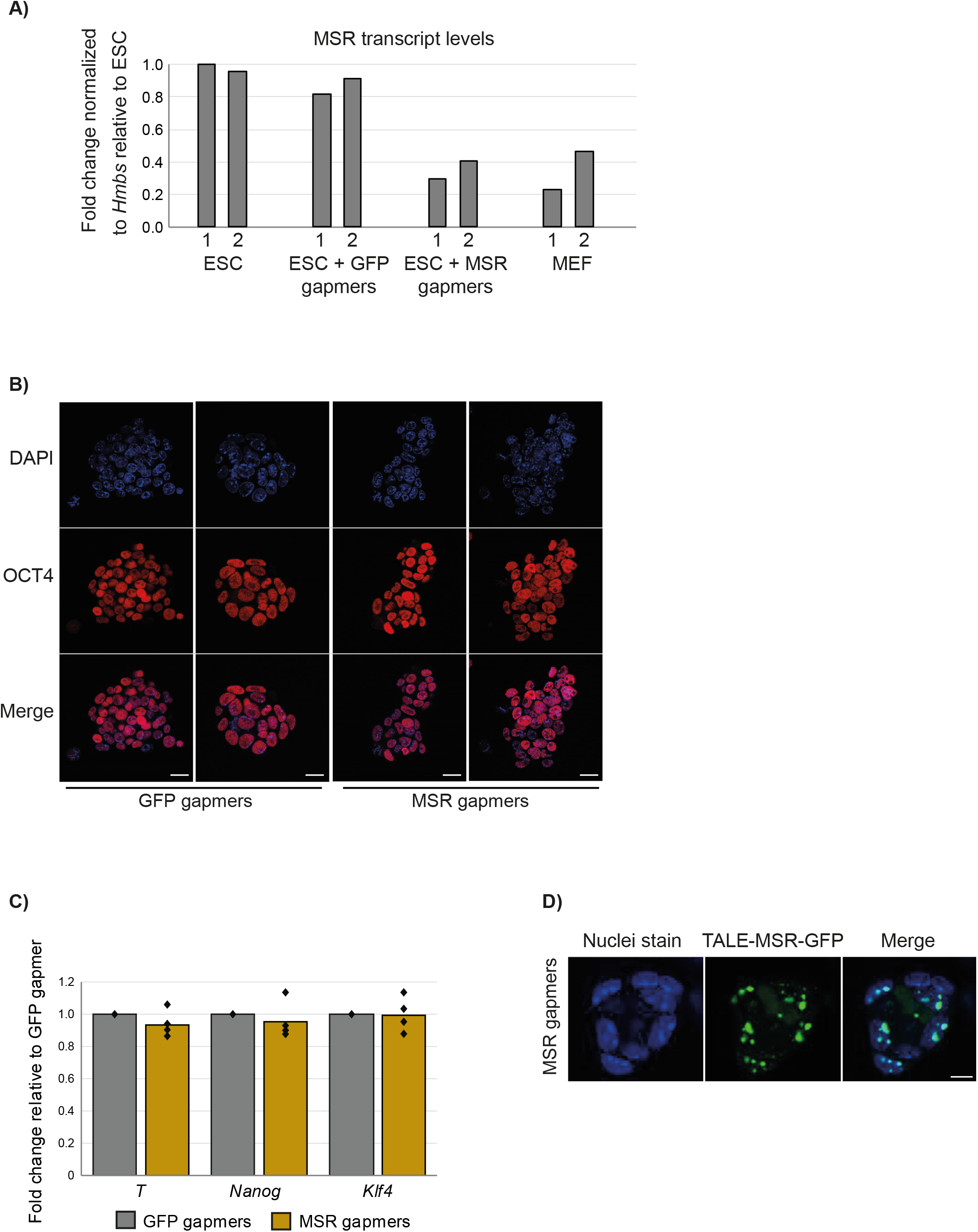
Embryonic stem cells do not differentiate when MSR transcript levels are reduced, Related to Figure 2. A) Histogram of MSR transcripts levels in ESCs that were either untransfected or transfected with GFP or MSR gapmers, and in mouse embryonic fibroblasts (MEFs). Expression levels are shown as fold-change relative to *Hmbs*. Two independent biological replicates are shown. B) Immunofluorescence microscopy images of OCT4 in ESCs, following transfection with GFP or MSR gapmers. DNA was counterstained with DAPI. Scale bars, 20μm. C) Histogram showing the transcript levels of undifferentiated ESC marker genes (*Nanog*; *Klf4*) and differentiated cells (*T*), as measured by RT-qPCR. Data were acquired from three independent experiments and each dot represents individual biological replicates. D) Representative images of live ESC nuclei showing TALE-MSR-GFP localisation at chromocenters is retained following transfection with MSR gapmers. Scale bar, 10μm. See also Supplementary Video 4.

**Supplementary Figure 3:**
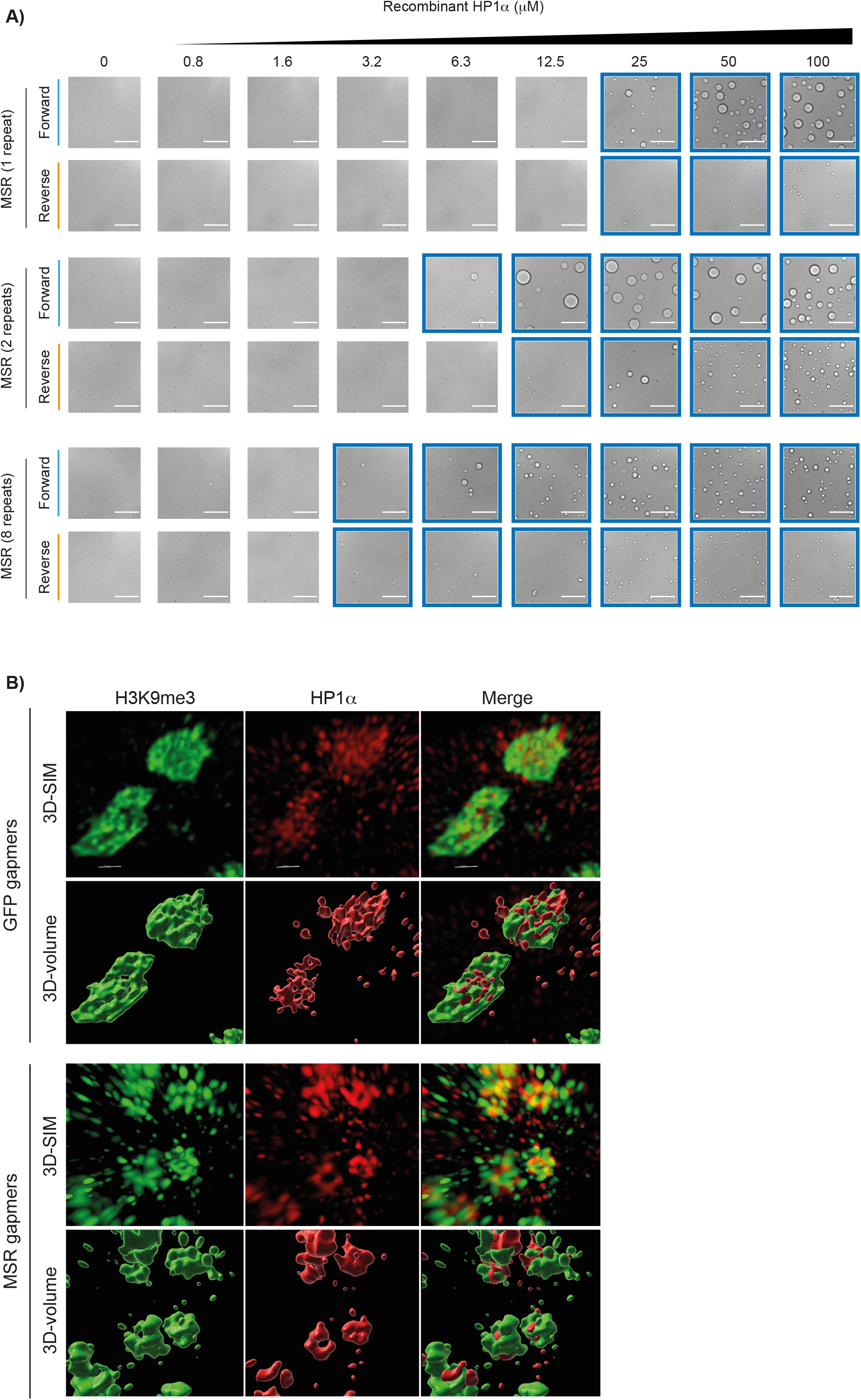
Satellite RNA promotes phase-separation of the heterochromatin protein HP1α, Related to Figure 3. A) Different lengths of forward MSR RNAs contribute to HP1α droplet formation *in vitro*. Images framed in blue show the conditions that allow for the formation of HP1α droplets. All images show the same magnification; scale bar, 100μm. B) Representative 3D-SIM images after LNA-DNA gapmer transfection. For each experiment, the top panel shows the 3D-SIM reconstruction of H3K9me3 (red), HP1α (green) and merged, and the lower panel shows the equivalent projected volumes.

**Supplementary Video 1: Visualising chromocenters throughout the cell cycle, Related to Figure 1.** Time lapse imaging of ESCs after doxycycline-induced expression of TALE-GFP against major satellite repeats (MSR). Images were acquired every 15 min over 72 time frames (18 hrs).

**Supplementary Video 2: Rapid movement of chromocenters, Related to Figure 1.** Time lapse imaging (compiled five frames per second) of a representative ESC expressing doxycycline-induced TALE-MSR-GFP. Images were acquired every 2 min for 60 frames (2h).

**Supplementary Video 3: FRAP analysis of chromocenters, Related to Figure 1.** Time lapse imaging (compiled five frames per second) of a representative ESC expressing doxycycline-induced TALE-MSR-GFP and after GFP gapmer transfection. One chromocenter was photobleached and images acquired every second for 310 seconds.

**Supplementary Video 4: FRAP analysis of chromocenters after MSR RNA depletion, Related to Figure 2.** Time lapse imaging (compiled five frames per second) of a representative ESC expressing doxycycline-induced TALE-MSR-GFP and after MSR gapmer transfection. One chromocenter was photobleached and images acquired every second for 310 seconds.

**Supplementary Table 1: Additional information related to the Online Methods.**

## Notes

### Competing Interest Statement

The authors have declared no competing interest.

